# SpoIIE drives asymmetric cell division in *B. subtilis* by sequential modulation of the cytokinesis machinery

**DOI:** 10.1101/2025.06.09.658746

**Authors:** Alexis Ryan, Georgia R. Squyres, Matthew J. Holmes, Alex Bisson, Ethan C. Garner, Niels Bradshaw

## Abstract

To form a dormant spore, *Bacillus subtilis* and related endospore-forming bacteria divide asymmetrically to generate daughter cells of unequal size, the smaller of which becomes the spore. The transmembrane protein SpoIIE repositions the cell division machinery and controls cytokinesis during sporulation, but the molecular basis for the precise placement of the asymmetrical division site to the quarter cell point is unknown. Here, we applied live-cell fluorescence microscopy techniques to reveal that SpoIIE localizes with the treadmilling components of the cell division machinery. We found that SpoIIE opposes the inhibitory activity of the MinCD complex, which prevents assembly of Z-rings near the cell poles. Cells expressing a variant of SpoIIE with its transmembrane region replaced by an unrelated transmembrane anchor assembled condensed Z-rings that were unable to initiate constriction. This reveals a new function of SpoIIE and a possible checkpoint licensing cytokinesis downstream of Z-ring condensation. Potentially explaining the role of SpoIIE in cytokinesis, we demonstrated that SpoIIE’s transmembrane region interacts with DivIB, an enigmatic structural component of the cell wall synthesis complex required for cytokinesis during sporulation. Finally, we found that FtsZ filaments are unusually short during sporulation, which requires the transmembrane domain of SpoIIE. Together, these results demonstrate that SpoIIE sequentially influences polar divisome assembly at distinct steps to drive asymmetric cell division.

## Introduction

How genetically identical cell populations differentiate to distinct cell types is a fundamental problem in developmental biology (Horvitz and Herskowitz 1992). A recurrent pathway to differentiation is to break symmetry through unequal partitioning of cellular factors between daughter cells following cell division (Kysela et al. 2012; Sunchu and Cabernard 2020). Sporulation in *Bacillus subtilis* is an established model system for addressing mechanisms of bacterial cell fate determination(Stragier and Losick 1996; Barak et al. 2019). The initial symmetry-breaking step of spore formation is a shift of cytokinesis from the midcell position to an asymmetric site near one cell pole. This yields a smaller cell that becomes a dormant spore and a larger cell that engulfs the forming spore and builds a protective coat around it. Despite decades of study, the molecular basis for orchestrating asymmetric cell division is elusive.

Bacterial cells exquisitely control the position and timing of cell division so that each daughter cell inherits the appropriate complement of biomolecules(Adams and Errington 2009; Reyes-Lamothe and Sherratt 2019). Cytokinesis is directed by localized assembly of a conserved complex of dynamic proteins called the divisome (Du and Lutkenhaus 2017; McQuillen and Xiao 2020; Barrows and Goley 2020). The timing of cytokinesis is limited by the recruitment of cell wall synthesis enzymes to the divisome, but how these enzymes are recruited and whether there is a post-recruitment checkpoint before initiation of cell wall synthesis is not understood (Monteiro et al. 2018; Squyres et al. 2021; Yang et al. 2021; Perez et al. 2024; Schäper et al. 2024). While the particular mechanisms that control cell division are as diverse as bacterial species, many core components are widely conserved(Adams and Errington 2009; Eswara and Ramamurthi 2015; Barrows and Goley 2020).

Whereas vegetatively growing *B. subtilis* divides medially to produce identical daughter cells in nutrient-rich environments, starvation initiates a developmental program that repositions the division septum near one cell pole to generate daughter cells of different sizes (Stragier and Losick 1996; Barak et al. 2019). The small cell resulting from this asymmetric division becomes the dormant spore. Cytokinesis during endospore formation is additionally modified to produce an unusually thin septum(Barák and Youngman 1996; Khanna et al. 2021). The sporulation-specific transmembrane protein SpoIIE directs efficient polar placement of the division septum and production of thin septal cell wall, but the molecular mechanisms by which SpoIIE executes these functions have remained mysterious(Barak et al. 2019). Here, we use a combination of genetics and live-cell single molecule microscopy(Bisson-Filho et al. 2017; Squyres et al. 2021) to uncover how SpoIIE controls divisome placement and cytokinesis.

One clue to how SpoIIE controls cell division is that it colocalizes with the divisome(Arigoni et al. 1995; Levin et al. 1997). The divisome is composed of two treadmilling polymeric proteins, FtsZ, a GTPase related to tubulin, and FtsA, a peripherally membrane associated ATPase related to actin(Bisson-Filho et al. 2017; Yang et al. 2017) (Fig. 1A). FtsZ additionally interacts with a set of bundling proteins that are important for the assembly and condensation of the divisome, collectively referred to as Z-ring Binding Proteins (ZBPs) (Squyres et al. 2021)(Fig. 1B). While FtsA and FtsZ are nearly universally conserved, there is more diversity in the makeup of ZBPs across bacterial phyla(Eswara and Ramamurthi 2015). These proteins assemble a dynamic treadmilling ring that moves circumferentially, perpendicular to the long axis of the cell(Bisson-Filho et al. 2017; Yang et al. 2017). It is attractive that SpoIIE could interact with the divisome similar to a ZBP because FtsZ treadmilling and FtsA/Z filament bundling are critical for positioning the septum and controlling the initiation of cytokinesis(Yang et al. 2021; Whitley et al. 2021), and SpoIIE localization depends on FtsZ(Levin 1997). However, there is uncertainty as to how SpoIIE interacts with the divisome. Although SpoIIE constricts along with the divisome, (Bradshaw and Losick 2015) its localization appears shifted towards the pole compared to FtsZ(Eswaramoorthy et al. 2014; Khanna et al. 2021; Chareyre et al. 2024).

**Figure 1:**
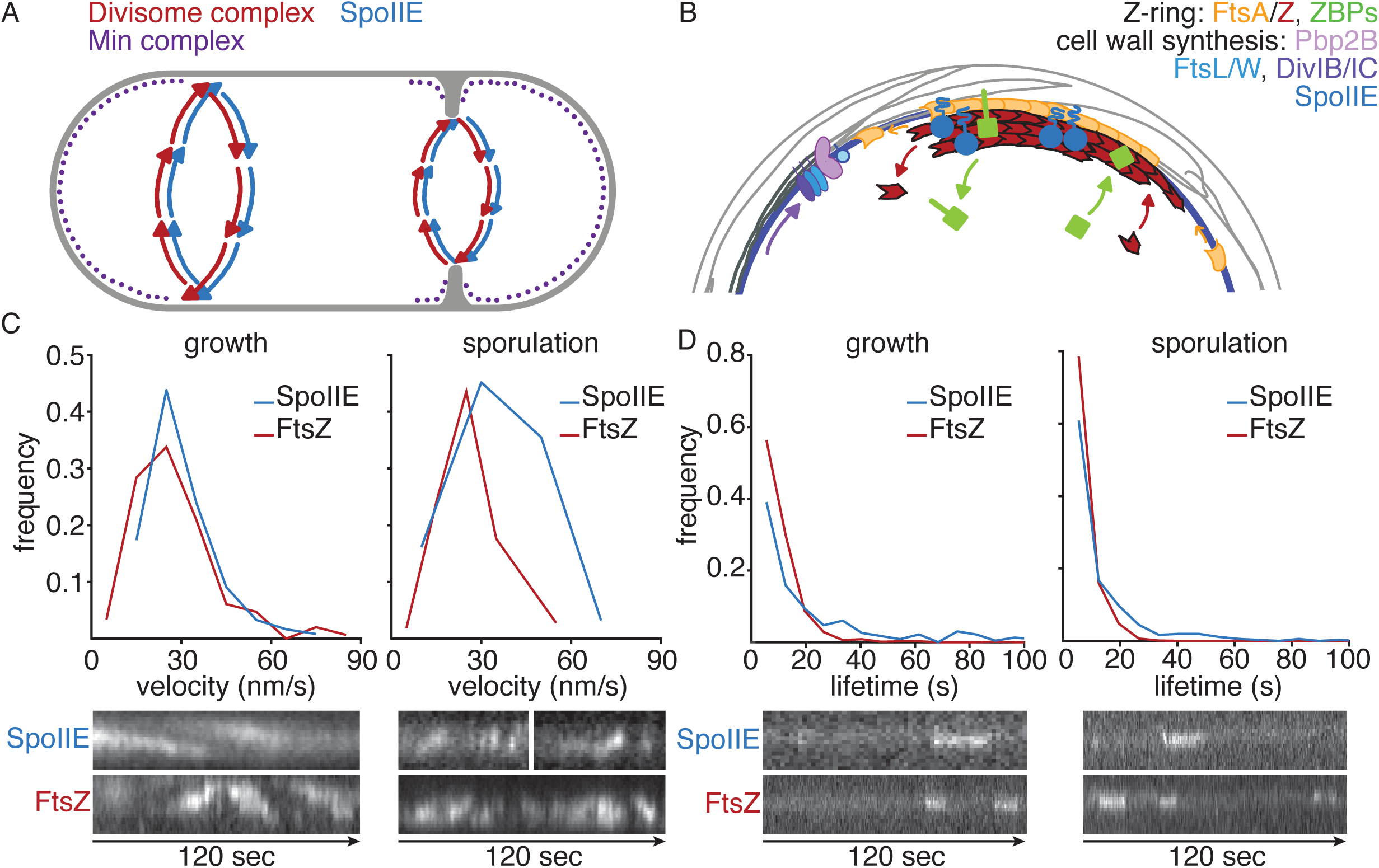
SpoIIE associates with the directionally moving divisome complex. **(A)** A cartoon depicting the localization of the divisome (red), SpoIIE (blue), MinC/D (purple) in a longitudinal cross section of a sporulating cell is shown. The divisome on the right is depicted as undergoing constriction as indicated by invaginating septal cell wall (grey). The divisome and SpoIIE are indicated with arrows to indicate that they are undergoing circumferential motion around the cell. **(B)** A cartoon depicting the divisome complex is shown in a half lateral cross section of a cell. Treadmilling filaments of FtsA (orange) and FtsZ (red) are shown anchored to the cell membrane (blue) by FtsA. Molecules of FtsZ-associated ZBPs (ZapA, SepF, EzrA; green) bind to stationary FtsZ subunits, while cell wall synthesis regulators (Pbp2B, DivIB/IC, FtsL/W; purple) move directionally with a similar velocity to each other, depositing new septal cell wall material (gray). Membrane-bound SpoIIE (blue) is shown colocalized with the divisome, and single molecules are stationary like a ZBP. **(C)** Plots are shown of the bulk velocity distribution of FtsZ-Halo (red) and SpoIIE-Halo (blue) ectopically expressed during vegetative growth (left) or under sporulation conditions (right). Velocities were calculated from kymographs like those shown below the plots. Slopes of tracks across the cell axis over time are used to quantify velocity of bulk protein movement. Images are 1.2uM in height. **(D)** Plots of single molecule lifetime distributions of FtsZ-Halo (red) and SpoIIE-Halo (blue) are shown. Lifetimes were measured using TrackMate single particle tracking with sample traces shown below. Images are 2.5um in height.

A principal mechanism that controls divisome positioning in medially dividing cells is executed by the proteins MinC and MinD, which inhibit FtsZ assembly at polar sites(Gregory et al. 2008; Szwedziak and Ghosal 2017). In endospore forming bacteria, MinC/D are concentrated near the cell poles, at the edges of constricting divisomes, and near recently disassembled divisome rings by interaction with the curvature sensing protein DivIVA (mediated by MinJ) (Ramamurthi and Losick 2009; Eswaramoorthy et al. 2014; Chareyre et al. 2024)(Fig 1A). While MinC/D are still present during asymmetric division in sporulation, the mechanism by which SpoIIE overcomes MinC/D activity at polar sites is not known(Barak 2014; Barak and Muchová 2018). A secondary mechanism to control divisome positioning, nucleoid occlusion, is also present in *B. subtilis* and is counteracted by a sporulation specific factor RefZ, which fine tunes divisome positioning and creates some redundancy with SpoIIE in promoting asymmetric division during sporulation(Wagner-Herman et al. 2012; Miller and Herman 2022; Muchová et al. 2024).

For cytokinesis to occur, the divisome undergoes condensation dependent on the ZBPs, which coincides with recruitment of a complex of proteins that direct cell wall synthesis(Squyres et al. 2021; Whitley et al. 2021). In *B. subtilis* this complex (which we refer to as the “divisome core complex” (Goley et al. 2011; Käshammer et al. 2023), but which has also been termed the “outer ring” (Daniel et al. 2006)) consists of Pbp2B, FtsL, FtsW, DivIB, and DivIC. Cell wall synthesis powers circumferential movement of these proteins, but the mechanisms that initiate and specify the thickness of septal peptidoglycan are not known(Squyres et al. 2021; Yang et al. 2021; McCausland et al. 2021; Whitley et al. 2021; Schäper et al. 2024). During sporulation, cytokinesis is modified by SpoIIE to produce unusually thin septa. FtsA/Z filaments are shifted away from the cell pole at the leading edge of the constricting septum, while SpoIIE is localized towards the pole (Khanna 2021,Eswaramoorthy 2014). Importantly, these roles of SpoIIE in cytokinesis can be genetically separated from the role of SpoIIE in assembling divisomes at polar sites; replacement of the native transmembrane domain of SpoIIE with two unrelated transmembrane segments from *E. coli* MalF allows assembly of polar divisome rings that produce thick septa and have symmetrically distributed FtsA/Z when they constrict(Khanna 2021). The pole-shifted distribution of SpoIIE during cytokinesis additionally depends on DivIVA, linking SpoIIE localization to spatial cues in the cell(Eswaramoorthy et al. 2014; Chareyre et al. 2024). However, the mechanism by which SpoIIE shapes the septum and the importance of asymmetric FtsA/Z localization has not been established.

Here, we applied live-cell TIRF microscopy to determine how SpoIIE interacts with and modulates the cell division machinery. We find that SpoIIE acts like a specialized ZBP that opposes MinC/D, suggesting a mechanism for how SpoIIE dictates the position of the polar divisome. Additionally, we find that SpoIIE has a second function to initiate cytokinesis following divisome condensation. Correlated with this function, SpoIIE shortens FtsZ filaments and interacts with DivIB. This provides a mechanistic framework for how polar division is orchestrated during endospore formation, reveals a role for SpoIIE recruiting the divisome core complex and initiating cytokinesis, and identifies SpoIIE as a modulator of FtsZ dynamics.

## Results

### SpoIIE associates with the directionally moving divisome complex

Although SpoIIE colocalizes with Z-rings, it was not known whether SpoIIE associates with treadmilling FtsZ (as do ZBPs), moves with the cell wall synthesis complex, or localizes near divisomes (like DivIVA) (Eswaramoorthy et al. 2014; Bradshaw et al. 2017; Barak et al. 2019; Chareyre et al. 2024)(Fig. 1B). First, we assayed how SpoIIE moves at the Z ring, with the expectation that fully labeled SpoIIE would appear to move directionally with the same velocity as the underlying treadmilling FtsZ filaments (Squyres et al. 2021). We ectopically expressed *spoIIE-halo* during vegetative growth, labeled cells with JF549, and recorded time lapse movies by TIRF microscopy of SpoIIE. We found that the bulk signal of fully-labeled SpoIIE appears to move directionally with a velocity of 29 nm/sec, indistinguishable from our measured FtsZ velocity (Fig. 1C) or other characterized divisome proteins (Bisson-Filho et al. 2017; Squyres et al. 2021). Though high levels of SpoIIE expression make velocities challenging to quantify for *spoIIE-halo* expressed under its native promoter during sporulation, we observed similar apparent bulk directional movement of densely labeled SpoIIE at sporulation divisomes (Fig. 1C). Thus, SpoIIE associates with the division complex independent of sporulation-specific factors.

Z-ring binding proteins (ZBPs), like ZapA, SepF, and EzrA, directly bind to FtsZ filaments and, like FtsZ monomers, appear stationary at a single molecule level. In contrast, single molecules of proteins associated with the divisome core complex move directionally with similar velocities to each other (Squyres et al. 2021; Schäper et al. 2024). To determine which divisome subcomplex SpoIIE is associated with, we imaged single molecules of SpoIIE-Halo at divisomes during sporulation. We found that SpoIIE molecules are stationary both at polar divisomes during sporulation and at medial divisomes during growth (Fig. 1D). These molecules persist at divisomes with an average lifetime of 12 seconds during sporulation and 24 seconds during growth (Fig. 1D). Notably, these lifetimes are longer than the lifetime of FtsZ subunits (6.5 seconds at divisomes during sporulation and 9.5 seconds at divisomes during growth; Fig. 1D)(Bisson-Filho et al. 2017; Squyres et al. 2021), suggesting that SpoIIE either interacts with multiple FtsZ filaments or does not diffuse away once FtsZ treadmills away. Thus, SpoIIE is part of the divisome complex and directly or indirectly binds to FtsZ filaments similar to the FtsZ bundling factors ZapA, SepF, and EzrA (Squyres et al. 2021). However, SpoIIE molecules persist at the division site for longer than the underlying FtsZ subunits.

To determine whether the immobile ZBP-like behavior of SpoIIE correlates with its functions for divisome positioning and/or cytokinesis, we genetically separated these roles by replacing the SpoIIE transmembrane domain with two transmembrane segments from *E. coli* MalF (MalF-SpoIIE^cyt^)(King et al. 1999; Carniol et al. 2005). Strains expressing *malF-spoIIE^cyt^* fail to produce thin septal peptidoglycan, and FtsZ is symmetrically localized at the septum rather than biased toward the mother side of the cell(Khanna et al. 2021) (Fig. 2A,B). Repeating TIRF-microscopy assays with *malF-spoIIE^cyt^-halo* expressed in growing cells demonstrated that the bulk velocity of fully labeled MalF-SpoIIE^cyt^ at divisomes (23nm/s) was the same as SpoIIE and FtsZ (29nm/s and 27 nm/s, respectively)(Fig. 2C), and single molecules were stationary. Thus, this fragment of SpoIIE binds to FtsZ filaments like a ZBP and is sufficient to reposition the septum. We note that the single molecule lifetime of MalF-SpoIIE^cyt^ was reduced to 9.1 seconds, still longer than the lifetime of FtsZ at these asymmetric division sites, but shorter than that of full-length SpoIIE (Fig. 2D). Thus, SpoIIE acts as a sporulation-specific ZBP that repositions the divisome for polar division.

**Figure 2:**
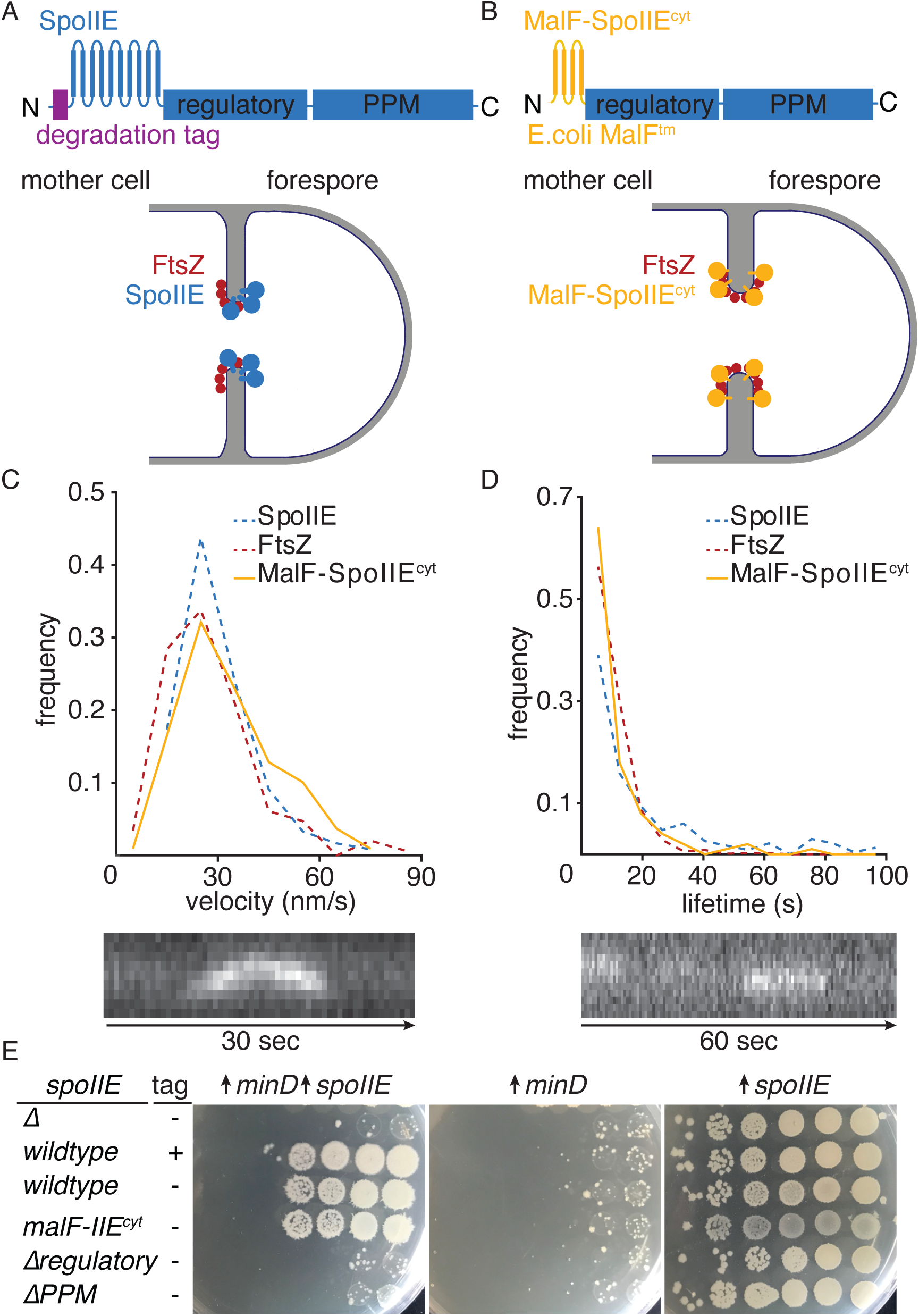
SpoIIE counteracts MinCD to promote asymmetric divisome assembly. **(A)** A cartoon depicting the domain architecture of SpoIIE is shown with N-terminal degradation tag, multipass transmembrane region, regulatory region, and PPM family serine/threonine phosphatase domain indicated. Below is a cartoon depicting SpoIIE (blue) and FtsZ (red) localization in a lateral cross section of a cell with the invaginating cell wall shown in grey. **(B)** A domain diagram depicts the MalF-SpoIIE^cyt^ construct (orange) in which the transmembrane segment of SpoIIE has been replaced with two transmembrane segments of *E. coli* MalF and the FtsH degradation tag (purple) has been deleted. As depicted in the cartoon below, while SpoIIE is distributed towards the forespore (right) MalF-SpoIIE^cyt^ is symmetrically localized. **(C)** A plot of velocity distributions of bulk labeled MalF-SpoIIE^cyt^-Halo (orange) expressed in vegetatively growing cells is shown with dashed lines depicting the distributions of FtsZ-Halo (red) and SpoIIE-Halo (blue) from figure 1 shown as a reference. Image of a representative kymograph from which the velocities were calculated is shown below (image height 1.4um). **(D)** A plot of lifetime distributions of single molecules of MalF-SpoIIEcyt-Halo (orange) expressed in vegetatively growing cell is shown as in C. A sample kymograph of stationary MalF-SpoIIE^cyt^ single molecules is shown below (image height 1.9um). **(E)** Plates showing growth of *B. subtilis* cells expressing various *spoIIE* constructs and/or overexpressing *minD*.

### SpoIIE is sufficient to oppose MinC inhibition of division

While MinC/D play an important role in restricting vegetative divisomes to the midcell, these proteins also contribute to polar divisome placement during sporulation(Jamroškovič et al. 2012; Barak and Muchová 2018). Whether the activities of SpoIIE and the Min complex intersect is not known. Intriguingly, ZapA, a vegetative ZBP, was discovered on the basis of its ability to counteract MinCD when overexpressed(Gueiros-Filho and Losick 2002). Thus, we hypothesized that SpoIIE might similarly oppose the activity of the Min complex and that this could contribute to polar divisome positioning during sporulation. To test this, we overexpressed *minD* in vegetative cells, which relocalizes activated MinC throughout the cell thereby inhibiting Z-ring assembly, killing cells(Gueiros-Filho and Losick 2002). We then assayed whether ectopically expressed *spoIIE* restores growth under these conditions. While *malF-spoIIE^cyt^* was sufficient to rescue growth when MinD was overexpressed, expression of *spoIIE* constructs lacking portions of the cytoplasmic domain that are required for divisome colocalization (Carniol et al. 2005) cannot rescue growth (Fig. 2E). These results indicate that SpoIIE opposes the Min complex and that this activity correlates with the interaction between SpoIIE and treadmilling FtsZ filaments.

### SpoIIE alters FtsZ polymerization during sporulation

It has long been hypothesized that SpoIIE directly or indirectly influences FtsZ as a part of its role promoting asymmetric cell division. To address this, we first asked whether deletion of *spoIIE* altered FtsZ treadmilling. FtsZ velocity at divisomes was unchanged by deletion of *spoIIE* during sporulation (wildtype 25 nm/s; *ΔspoIIE* 28 nm/s) and was indistinguishable from FtsZ movement during vegetative growth (25 nm/s)(Bisson-Filho et al. 2017; Squyres et al. 2021) (Fig. 1C, 3A). However, deletion of *spoIIE* increased the single molecule lifetime of FtsZ at divisomes during sporulation by roughly 50% (mean lifetime of wildtype 6.5 s; *ΔspoIIE* 8.9 s), similar to the lifetime observed at vegetative divisomes (9.5 s) (Fig. 1C, 3B). Intriguingly, we obtained similar lifetimes for FtsZ particles at medial and polar divisomes in sporulating cells (Fig. S1). These effects were similar to the effect previously observed for EzrA, which sequesters FtsZ monomers from the overall pool in addition to its primary activity bundling FtsZ filaments (Squyres et al. 2021). The changes in filament length as we observe for deletion of *spoIIE* are similar to effects seen for *ezrA*, although *ezrA* deletion causes polar divisomes to assemble, while *spoIIE* deletion depresses polar division during sporulation (Levin et al. 1999). This led us to question whether FtsZ filament shortening correlates with the activity of SpoIIE to promote polar divisome assembly or to the role of SpoIIE during cytokinesis. Again, we genetically uncoupled these roles by expressing *malF-spoIIE^cyt^.* In contrast to our previous results that *malF-spoIIE^cyt^* is sufficient to counteract MinC/D (Fig. 2E), cells expressing *malF-spoIIE^cyt^* had FtsZ filaments of comparable length to *ΔspoIIE* cells, despite having higher protein levels than SpoIIE (23 nm/s velocity, 9.0 s mean lifetime) (Fig. 3A, B). This suggests that MalF-SpoIIE^cyt^ is unable to sequester FtsZ monomers in a manner that results in FtsZ filament shortening, which is not required to asymmetrically position the divisome. While it is not clear whether the length of FtsZ filaments is important for cytokinesis during sporulation, these results establish SpoIIE as a modulator of FtsZ polymerization, potentially by binding and sequestering FtsZ monomers.

**Figure 3:**
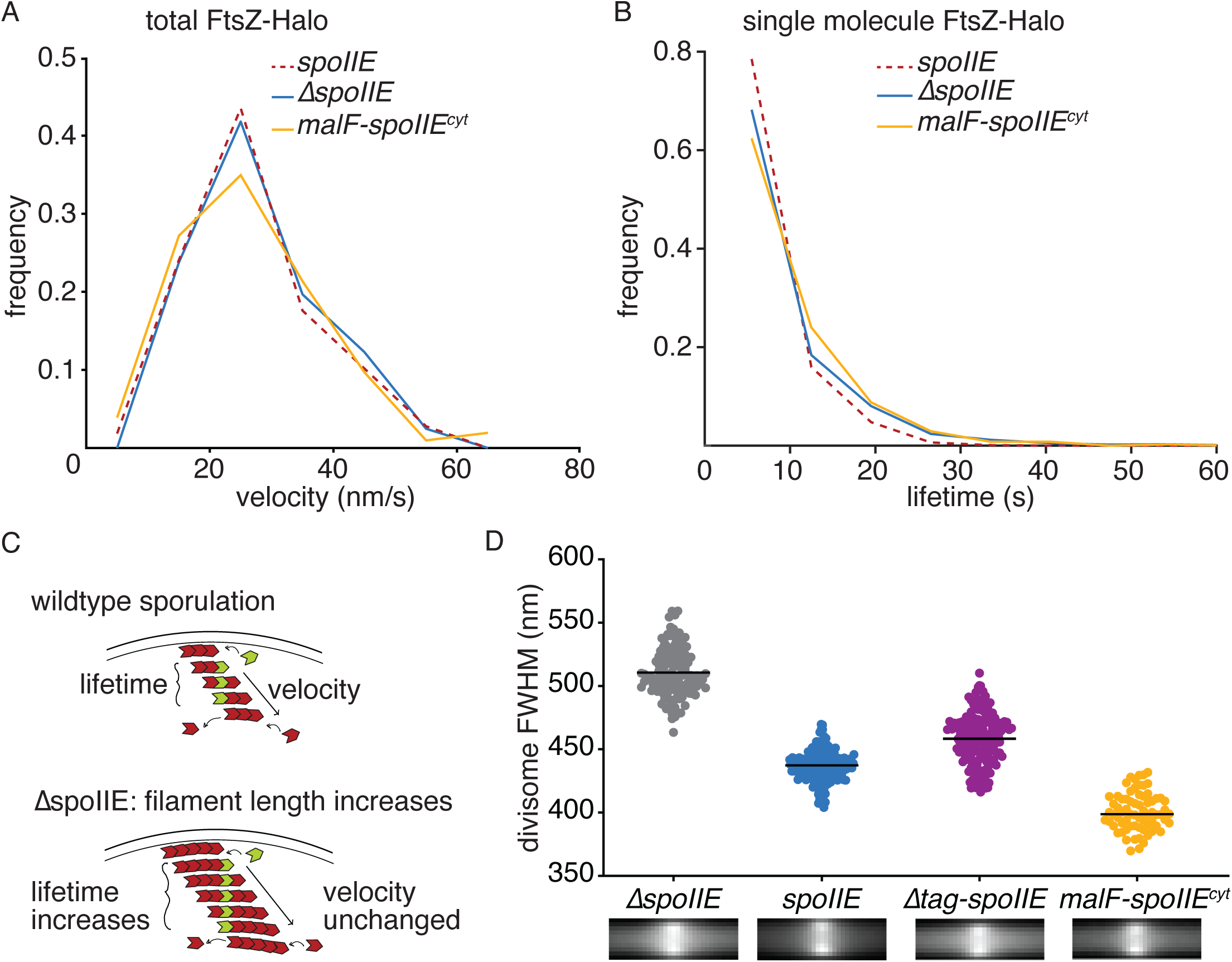
SpoIIE shortens FtsZ filaments during sporulation. **(A)** Plots of velocity distributions of bulk labeled FtsZ-Halo in wildtype cells (dashed red line, data reproduced from Figure 1), *ΔspoIIE* cells (blue), and *malF-spoIIE^cyt^*cells (orange) during sporulation. **(B)** Plots of lifetime distributions of single molecules of Halo-tagged FtsZ from strains as in A. **(C)** A cartoon of the relationship between filament length and treadmilling velocity with a single subunit of a treadmilling FtsZ subunit shown in green. When velocity remains the same but single molecule lifetime increases, the length of filaments can be extrapolated to have increased. **(D)** A violin plot of the full width half maximum (FWHM) measurements of individual divisomes from sporulating cells with bar marking mean FWHM for cells sporulated with indicated SpoIIE constructs. Corresponding sum projections of all Z-rings for each condition pictured below data points. n > 75 cells each condition.

### SpoIIE is promotes divisome constriction after condensation

Following assembly of a divisome, two events are required to initiate constriction: condensation of the divisome ring (mediated by the ZBPs), followed by recruitment and activation of the cell wall synthesis machinery(Squyres et al. 2021; Whitley et al. 2021; Schäper et al. 2024). We therefore imaged divisomes in sporulating cells expressing variants of *spoIIE*, and determined whether condensation occurred by calculating the width of the average divisome projection (Fig. 3D). We used cells deleted for *σ^F^*, which blocks further development after asymmetric division making it easier to detect cell division events. We observed that deletion of *spoIIE* results in wider Z-rings (511 nm FWHM) than when wildtype *spoIIE* (437 nm FWHM) or *Δtag-spoIIE* (458nm) is expressed (Fig. 3D). This demonstrates that SpoIIE promotes Z-ring condensation, consistent with its role in promoting cytokinesis. Strikingly, Z-rings of *malF-spoIIE^cyt^* expressing cells were hypercondensed (401 nm FWHM) (Fig. 3D), suggesting that condensation occurred. Consistent with this, we found that *malF-spoIIE^cyt^*cells have a substantial reduction in the frequency with which they initiate cytokinesis judged by the membrane-intercolating dye FM4-64 (Fig. 4A,B). One third fewer divisomes of cells expressing *malF-spoIIE^cyt^* initiated cytokinesis compared to *Δtag-spoIIE* cells (41% vs. 63%) and ten times fewer cells were disporic (cells that have divided at both polar sites) (3% vs. 30%). These data suggest that there is a significant delay between divisome condensation and initiation of cytokinesis in *malF-spoIIE^cyt^* cells, revealing a possible checkpoint between these events. (SUMMARY OF CONCLUSIONS OF THIS SECTION)

**Figure 4:**
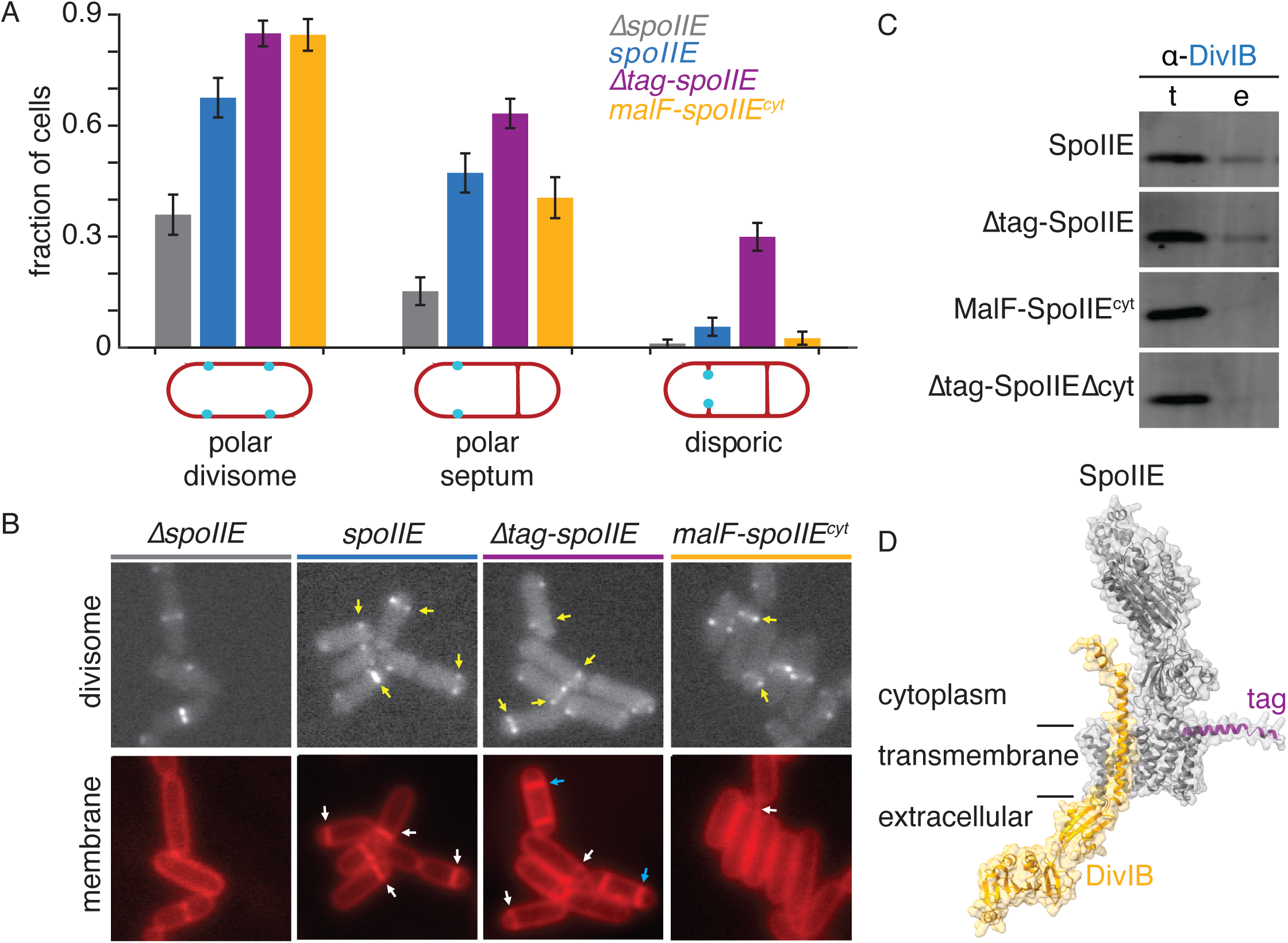
SpoIIE recruits DivIB and promotes divisome constriction. **(A)** Plots of quantification of fractions of cells with asymmetric Z-rings, septa, and disporic attempts for each SpoIIE construct represented in (A). Error bars are SEP with 95% confidence from n > 79 cells each condition. Cartoons are shown below depicting cells of each class with the membrane shown in red and the divisome shown in blue. **(C)** Representative images of fields of cells from each strain shown in (B) are shown. **(D)** An AlphaFold3 model of the complex between DivIB (orange) and SpoIIE (grey) is shown with the transmembrane domain and orientation noted. **(E)** A plot of the predicted alignment error (PAE) of the complex prediction is shown with the confidence of the prediction with a blue (0) yellow (10) white (30) color scheme. The transmembrane domain of each protein is indicated with a black bar above. **(F)** Western blots with an α-DivIB antibody are shown from the load (t) and elution (e) of co-immunoprecipitation of DivIB with the 3XFLAG-constructs of SpoIIE as indicated on the left.

### SpoIIE recruits DivIB to polar divisomes

Given that *malF-spoIIE^cyt^*is sufficient for *spoIIE* dependent Z-ring condensation, but not *spoIIE* dependent cytokinesis, we hypothesized that *malF-spoIIE^cyt^* cells do not properly recruit or activate the cell wall synthesis complex. We performed a pulldown and mass-spectrometry experiment to identify protein partners that interact with SpoIIE-3XFLAG. From this experiment we identified a component of the septal cell wall synthesis complex, DivIB, as the protein with the highest coverage and most unique peptides (Table S1). Additional pulldown experiments followed by western blots using Div-IB antibodies demonstrated that SpoIIE-3XFLAG efficiently pulled down DivIB, in contrast to MalF-SpoIIE^cyt^-3XFLAG or the transmembrane domain of SpoIIE alone, which did not (Fig. 4C, S2A). This raises the possibility that SpoIIE recruits DivIB (and perhaps the rest of the divisome core complex) to drive cell wall synthesis during sporulation.

Several additional lines of evidence support a role for SpoIIE interaction with DivIB in promoting cytokinesis during sporulation. First, while DivIB is non-essential for vegetative division (except at elevated temperatures), it is required for septation during sporulation, and the rare polar septa formed by *ΔdivIB* cells are thicker than wildtype sporulation, phenocopying *malF-spoIIE^cyt^*cells(Harry et al. 1993, 1994; Rowland et al. 1997; Katis and Wake 1999). Second, AlphaFold3 prediction of the complex between DivIB and SpoIIE yields a moderate-confidence model that places the transmembrane segment of DivIB in a conserved groove formed by helices α6, α7, and α9 of the SpoIIE transmembrane domain (Fig. 4D). Notably, in structural models of the divisome core complex, the transmembrane domain of DivIB is not associated with the rest of the complex and could be accessible for SpoIIE binding(Schäper et al. 2024).

## Discussion

How endospore-forming bacteria reposition the cell division site for sporulation is an enduring mystery(Barak 2019,Arigoni 1995,Carniol 2005,Ben-Yehuda 2002,Khanna 2021,Eswaramoorthy 2014). Here we report the discovery that SpoIIE, the sporulation-specific protein responsible for efficient asymmetric division, acts at sequential transitions to reposition and shape the division site. First, SpoIIE counteracts inhibition of polar division by MinC/D, correlated with its action as a sporulation-specific Z-ring-binding protein (ZBP) and suggesting a mechanism for how SpoIIE repositions the divisome to polar sites. Second, SpoIIE promotes condensation of the assembled Z-ring, consistent with its mechanistic similarities to previously characterized ZBPs. Third, SpoIIE recruits DivIB, and possibly the rest of the divisome core complex, to polar divisomes, suggesting a mechanism for how SpoIIE promotes initiation of cytokinesis and controls the thickness of septal cell wall. Together these events establish de-novo asymmetry in cell size that is translated into the cell-specific-transcriptional program that drives endospore development. These findings additionally raise several questions:

*How does SpoIIE counteract MinC/D?* MinC and MinD are broadly conserved across bacterial species and prevent assembly of divisomes near cell poles and adjacent to recent sites of division (Gregory et al. 2008; Szwedziak and Ghosal 2017). How SpoIIE overcomes this inhibition for programmed polar division during sporulation has been a longstanding question(Barak and Muchová 2018). One explanation for this could be that SpoIIE bundles FtsZ filaments similar to the ZBPs, which also counteract delocalized MinC/D when overexpressed (Gueiros-Filho and Losick 2002). Our finding that SpoIIE promotes divisome condensation (Fig. 3D), like the ZBPs(Squyres 2021) further supports this hypothesis. Complementary and intriguing possibilities are that SpoIIE could directly bind and neutralize MinC/D or occlude the MinC binding site(s) on FtsZ. Because *minC* deletion causes minicell formation, we hypothesize that SpoIIE activity to counteract MinC/D is a significant part of its mechanism of assembling polar divisomes.

*How does SpoIIE interact with the divisome?* Our finding that SpoIIE moves with indistinguishable velocity to treadmilling FtsZ filaments, while exhibiting stationary single-molecule behavior strongly suggests that SpoIIE directly interacts with the divisome similar to a ZBP. The previously characterized ZBPs (SepF, EzrA, and ZapA from *B. subtilis*, as well as others from other organisms) all directly interact with FtsZ (Squyres et al. 2021). However, despite one report of a direct interaction between SpoIIE and FtsZ in the literature (Lucet et al. 2000), we and others have not been able to detect such an interaction. Additionally, the longer lifetime of SpoIIE than FtsZ at divisomes (Fig. 1D, in contrast to the previously characterized ZBPs) suggests differences in how SpoIIE and the ZBPs interact with the Z-ring. Determining how SpoIIE interacts with the divisome will thus reveal how it exhibits sporulation specific ZBP activity.

*How does SpoIIE promote cytokinesis?* How cells transition from assembly of a treadmilling divisome to initiating cell-wall synthesis to drive cytokinesis is a major unanswered question (Squyres et al. 2021; Yang et al. 2021; Schäper et al. 2024). Previous studies of vegetative division in *B. subtilis* showed that ZBP-dependent condensation of the divisome coincides with initiation of cell-wall synthesis (Squyres et al. 2021; Whitley et al. 2021). Our results demonstrate that these steps can be uncoupled because cells expressing a fragment of SpoIIE that lacks the native transmembrane domain have hypercondensed divisomes that nonetheless fail to initiate constriction (Fig. 3D). One clue to this comes from our finding that SpoIIE interacts DivIB, an enigmatic member of the cell-wall synthesis complex that is required for sporulation (Fig. 4C). We hypothesize that SpoIIE interaction with DivIB helps to recruit and/or activate the cell wall synthesis complex under conditions that otherwise would be unfavorable to cytokinesis near the cell pole.

*How is SpoIIE phosphatase activity coordinated with cell division?* The end result of asymmetric cell division for sporulation is that SpoIIE becomes enriched in the small cell, where it initiates transcription from σ^F^ to begin the transformation of this cell into a spore(King et al. 1999; Bradshaw and Losick 2015; Chareyre et al. 2024). How SpoIIE is held inactive until cytokinesis is complete remains an important unanswered question. Our finding that SpoIIE interacts with the early treadmilling divisome complex containing FtsZ and the divisome core complex that drives cytokinesis suggests that these interaction partners could hold SpoIIE inactive. Additionally, it was recently discovered that SpoIIE interaction with DivIVA plays an important role localizing SpoIIE to the small cell and coordinating its cell-specific activation(Eswaramoorthy et al. 2014; Bradshaw and Losick 2015; Chareyre et al. 2024; Muchová et al. 2024). Future studies that reveal the molecular basis for these interactions will reveal how SpoIIE drives cell-fate determination that is coordinated with asymmetric cell division.

However, there is an apparent contradiction raised by the finding that SpoIIE interfaces with sequential components of the cell-division machinery. The treadmilling divisome assembly, the MinC/D positioning complex, the cell-wall synthesis regulator DivIB, and the curvature sensing protein DivIVA each exhibit unique localization and dynamic behavior, yet we only observe immobile molecules consistent with SpoIIE interacting with the FtsZ filaments similar to ZBPs. One possible answer to this is that SpoIIE may bring these factors together at the stage of divisome assembly to initiate the specialized asymmetric cell division required for endospore formation.

## Methods

### *B. subtilis* strains and molecular genetics

All *B. subtilis* strains were built in the PY79 background using natural competence (Harwood & Cutting 1990) and are listed in the table of strains. Cells were grown in Lennox lysogeny broth (LB, Sigma Aldrich) as liquid medium or as plates supplemented with 15% Bacto Agar (Difco). Plasmids constructed via isothermal assembly and validated by Sanger sequencing. Constructs integrated at native loci contained resistance markers and were patch streaked on antibiotics before stocking to verify insertion. All other constructs were inserted at the *amyE* or *lacA* loci and double crossover insertion of the desired gene was validated. Isothermal assembly products were verified by sequencing. All sporulation imaging strains used in this study were constructed lacking *σ^F^* (*ΔspoIIAC*) because progression of the sporulation program after cell division disassembles any existing Z-rings.

### Cell growth for imaging

Single colonies from plates freshly streaked from freezer stocks were grown to log phase at 25 °C overnight. For sporulation, cells were diluted into fresh 25% LB to achieve OD 0.025. Cells were grown on a roller at 37C until OD between 0.5-0.7. Cultures were pelleted and resuspended in an equal volume of defined sporulation resuspension media with fresh additives (Harwood and Cutting 2010). Sporulation imaging time points ranged from 1.5-3 hours as indicated (earlier for studying the divisome, later for studying septation). For vegetative growth cells were inoculated into fresh LB for epifluorescence imaging or CH media for TIRFm to achieve OD 0.0025 and grown at 37C. Cells were either assayed immediately during exponential growth (OD 0.2-0.3) or were back diluted in 1:100 series to be used throughout the day.

### HaloTag labeling

HaloTagged FtsZ was expressed from a second copy of *ftsAZ* under the pHyperSpank promoter at the *amyE* locus (Squyres et al. 2021). FtsZ was induced with 20 uM IPTG at the time of 100-fold back-dilution and incubated with JF549 or JF549x dye 1 hour before imaging. 1 nm dye was added for total labeling, and 40 pM dye was added for speckle labeling. Since SpoIIE is expressed at very high levels under its native promoter and is rapidly degraded by FtsH under vegetative expression, vegetative imaging of SpoIIE-Halo was initially conducted with saturating 0.5 mM IPTG. SpoIIE-Halo constructs for vegetative growth were either induced with 25 uM IPTG or 0.25% xylose, depending on strain background. Speckle labeling was achieved with 20pM JF549, and bulk labeling with 500pM JF549. FtsZ labeling conditions were identical during sporulation; however, HaloTagged SpoIIE constructs were instead expressed under the native SpoIIE promoter. Single molecules were imaged with 0.2 pM JF549, and bulk labeling with 100pM JF549.

### Epifluorescence microscopy

Images were acquired on an upright Olympus BX61 microscope scope using a 100X oil objective at room temperature. Cells were resuspended by flicking in 15 uL of sporulation resuspension media containing any relevant dyes (FM4-64 or DAPI), and 1 uL of sample was spotted onto a coverslip. A molded agarose pad made with sporulation resuspension media and 2% agarose was placed on top, and the assembly was placed on a slide with a custom drilled hole for air access to the sample.

### TIRF microscopy

TIRF movies and accompanying phase contrast and widefield fluorescence images were acquired on a custom Nikon Ti-E inverted microscope using a Nikon CFI Plan Apochromat DM Lambda 100X oil objective at 37C (Squyres et al. 2021). After spotting 2 uL of sample onto a glass-bottom MatTek dish, a pre-poured mould made with 2% agarose and CH media (or A+B media for sporulating cells) was placed on top. Samples were imaged at 37C in a heated chamber. Divisomes marked with ZapA-mNeonGreen were imaged with a 488 nm laser, and Halo constructs were imaged with a 561 nm laser.

Time lapse movies of FtsZ were recorded with 500 ms exposures for 2-4 minutes. Phase contrast and widefield fluorescence images were taken before and/or after time lapses. Total labeled movies to visualize velocities were recorded with an sCMOS camera, and single molecule movies were recorded using an emCCD camera. Since SpoIIE lifetimes were much longer than FtsZ lifetimes during vegetative growth, SpoIIE constructs were imaged ranging from 500 ms to 2 s frame rates. To ensure long-lived particles were single molecules, experiments were repeated varying laser power and exposure times to monitor photobleaching.

### Growth assays

Cultures were grown as described above until OD 0.4-0.5. Samples were pipetted into a sterile 96-well plate using water to bring all samples to OD 0.4. Wells were prepared with water to serially dilute cultures 10-fold out to 1:10,000. After pipetting to mix, 5 uL of each dilution series were spotted onto LB plates or LB plates containing each combination of inducers: 1 mM IPTG, 1% xylose, or 1 mM IPTG and 1% xylose. Replicates were grown for 14-16 hours at 37°C.

### Cell segmentation

For lifetime measurements and Z-ring width analyses, images were segmented using SuperSegger MATLAB code using the 100x *E. coli* preset identification parameters (Stylianidou et al. 2016). To identify sporulating cells and segment divisome images as well, the SuperSegger preset fluorescent parameters were used to segment ZapA-mNeonGreen and PspoIIE-mTurquoise2 images alongside phase contrast images.

### Single molecule lifetime measurements

Time lapse movies were read into ImageJ for particle tracking by TrackMate(Ershov et al. 2022). Spots of expected diameter 0.48 uM were detected with a manually set quality threshold to preliminarily link particle tracks from frame to frame. TrackMate files were exported to MATLAB and further analyzed using custom code as described(Squyres et al. 2021). After subtracting local background fluorescence, excluding overlapping molecules, and filtering out tracks not meeting a manually determined intensity threshold, tracks were fit to a Hidden Markov Model. Mean lifetimes were compared, and p-values were calculated with a Wilcoxon rank sum test. Significance of p < 0.05 was considered significant.

### Velocity measurements

Velocities from total labeled FtsZ movies were manually measured by kymograph analysis. A 5 pixel-wide line ROI was drawn across the short axis of cells in ImageJ and a kymograph drawn using the MultiKymograph plugin. Diagonal tracks representing movement across the cell axis over time were selected and their slopes measured.

### Quantification of Z-ring widths

Since the custom MATLAB code reads in Morphometrics-generated mask and pill mesh files, SuperSegger masks were run through Morphometrics to convert them. Cells masked as sporulating (containing PspoIIE-mTurquoise2 signal) were run through custom MATLAB code as described (Squyres et al. 2021) to measure fluorescence intensity of Z-rings (ZapA-mNeonGreen) and detect peaks. Z-ring full width half max measurements were calculated for each cell based fluorescence intensity from a 1 µm region surrounding each peak of ZapA-mNeonGreen intensity. The mean FWHM was calculated for cells of each genotype and sum Z-ring projections were generated from the aligned peaks.

### Asymmetric division rate quantifications

The same strains used for quantification of Z-ring widths were used to manually count fractions of cells with asymmetric Z-rings and/or septa. These strains are deleted for *σ^F^*, which means when they cannot progress beyond polar division in the sporulation program, they attempt to divide a second time at the other cell pole, resulting in a disporic cell with two septa. MicroManager epifluorescence image files were loaded into ImageJ. Using the Cell Counter plugin for ImageJ, cells were first counted for entry into sporulation cell division (CFP channel visible with a minimum fluorescence of 500 counts). These cells were then counted for the presence of two puncta representing medial or asymmetric Z-rings. The same cells were further sorted by septa. Cells were marked as having a medial septum, one polar septum, two polar septa, or no septa.

### Pulldowns and mass spectrometry

SpoIIE-3X was isolated from sporulating *B. subtilis* cells 2.5 hours post resuspension using anti-FLAG-M2-magentic beads (Sigma). Cells were lysed in a cell disruptor in 50 mM Tris-HCl, pH 8.5, 200 mM NaCl, supplemented with complete protease inhibitors (Roche). Membrane proteins were solubilized with 2% digitonin and extracts were incubated with anti-FLAG magnetic beads for 1 hour at 4°C. Samples were washed thoroughly in lysis buffer with 0.1% digitonin and eluted by competition with FLAG peptide for mass spectrometry or boiling in SDS buffer for westerns. Westerns were probed with anti-DivIB antibody provided as a generous gift form Elizabeth Harry. For mass spectrometry, a parallel sample from a strain expressing untagged *spoIIE* was used as a control. Each sample was run on an SDS PAGE gel and the lanes were cut into seven slices that were individually prepared for Mass-Spectrometry as marked on the gel image in Figure S3. Mass analysis was performed after trypsinization and elution from the gel by microcapillary reverse-phase HPLC, directly coupled to the nano-electrospray ionization source of an LTQ-Orbitrap Velos or LTQ-Oritrap XL mass spectrometer. These instruments are capable of acquiring individual sequence (MS/MS) spectra on-line at high mass accuracy (<2 ppm) and sensitivity (<<<1 femtomole) for multiple peptides in the chromatographic run. These MS/MS spectra are then correlated with known sequences using the algorithm Sequest (Eng et al. 1994)and programs developed in the Harvard University Mass-spectrometry laboratory(Chittum et al. 1998). MS/MS peptide sequences are then reviewed by a scientist for consensus with known proteins and the results confirmed for fidelity at the Harvard Mass Spectrometry and Proteomics Resource Laboratory, FAS Center for Systems Biology, Northwest Bldg Room B247, 52 Oxford St, Cambridge MA. Tabulated results are available as supplementary datasets.

## Acknowlegements

The authors thank Richard Losick and Bruce Goode for critical review of the manuscript and for input throughout the project. This research was supported by startup funds to NB from Brandeis University, the National Institute of General Medical Sciences of the National Institutes of Health award number 1R35GM158434, and by NSF MRSEC DMR-2011846. AR was supported by T32 GM007122. This work was supported by awards to AB: National Institute of General Medical Sciences of the National Institutes of Health under award number 1R35GM156992; the Human Frontiers Science Program (RGY0074/2021); National Science Foundation grant NSF-MBC2222076. AB is a Pew Scholar in the Biomedical Sciences, supported by The Pew Charitable Trusts.

---

**Figure S1:**
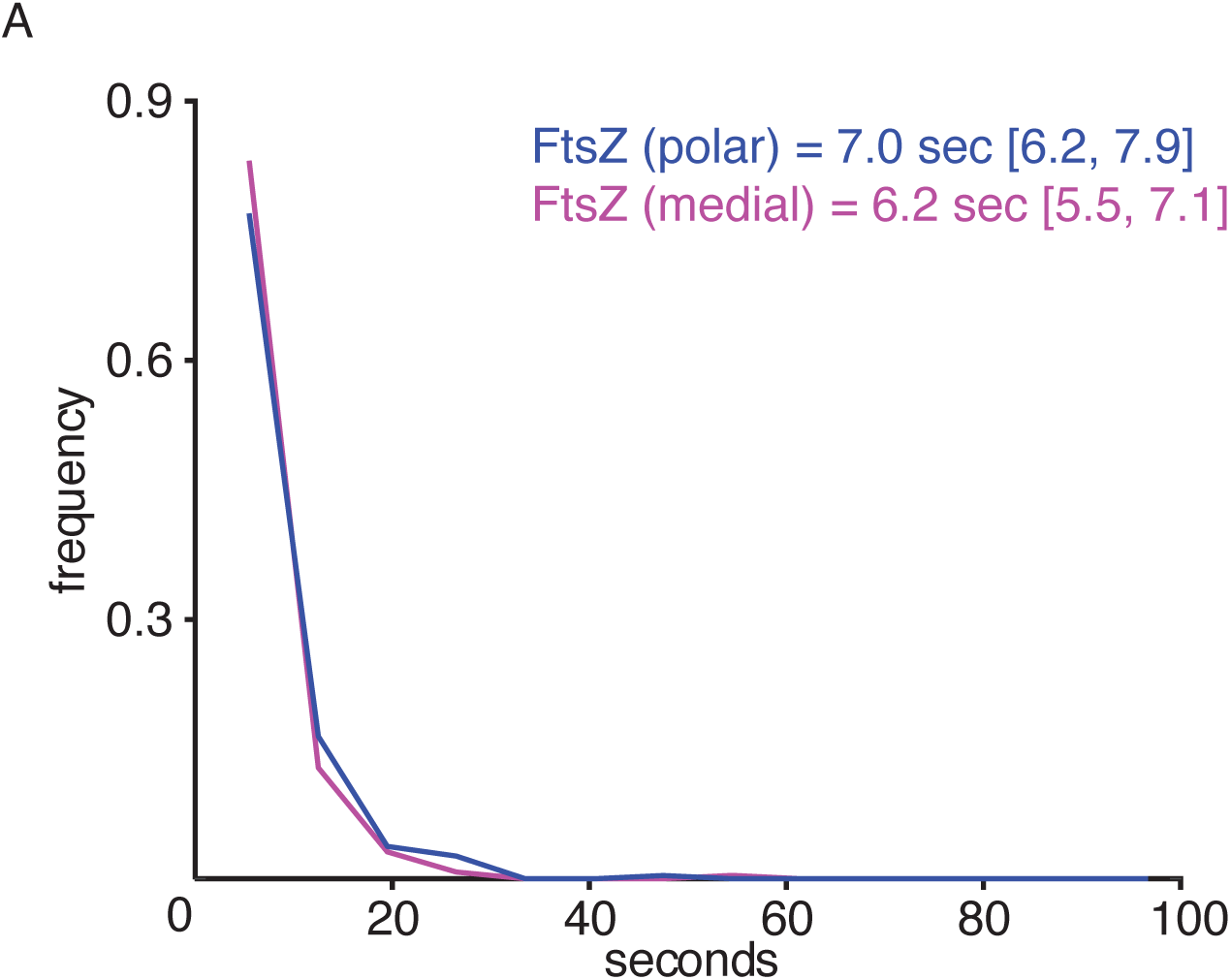
(A) Plot of single molecule lifetime distributions of FtsZ-Halo at medial (blue) and polar (purple) divisomes during sporulation are shown. Wilcoxon rank sum test concludes no significant difference (p = 0.4, n > 230 particles each).

**Figure S2:**
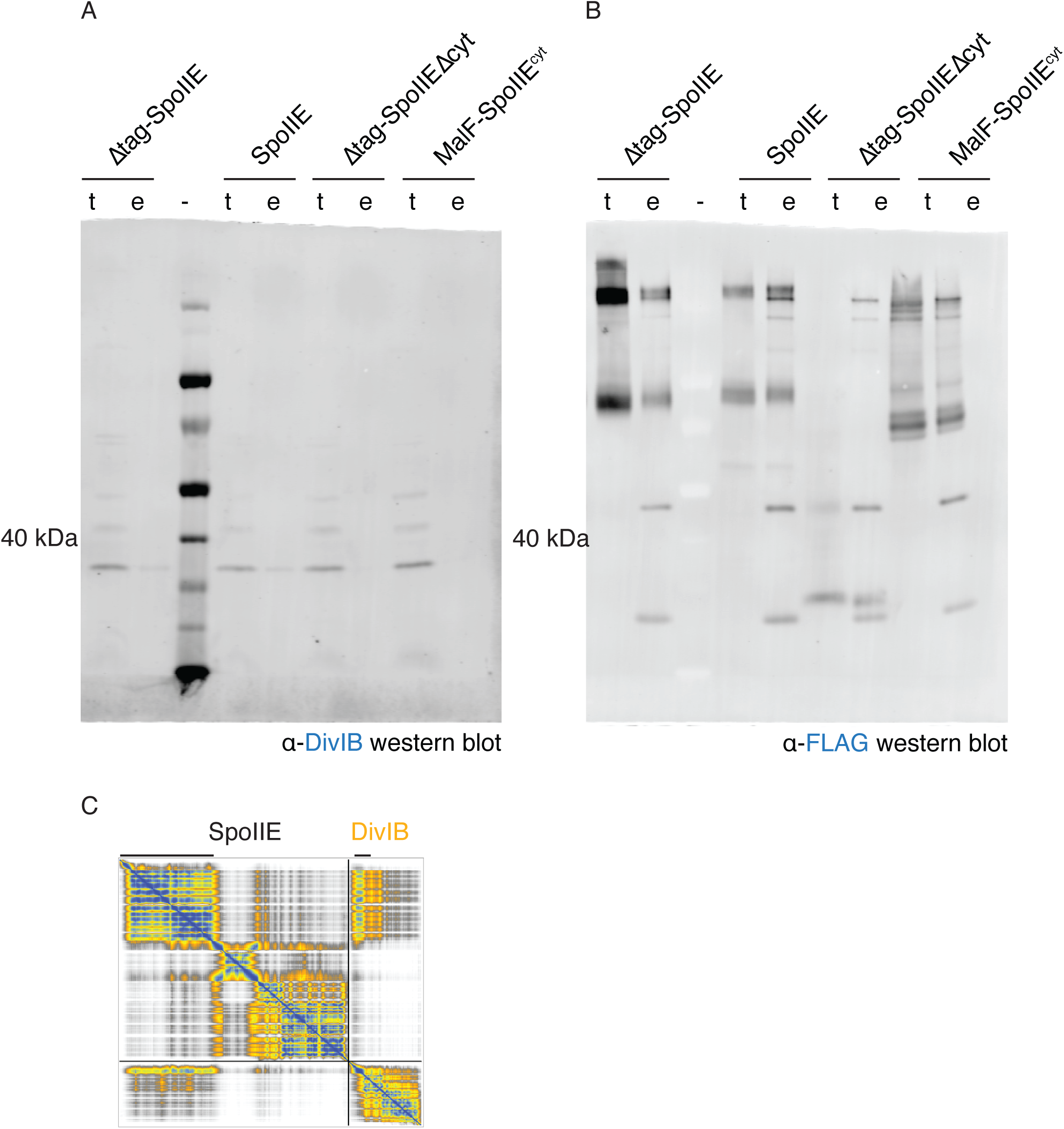
Western blots with (A) α-DivIB antibody and (B) α-SpoIIE antibody are shown from the load (t) and elution (e) of co-immunoprecipitation of DivIB with the 3XFLAG-constructs of SpoIIE as above the lanes. (C) Epifluorescence images at 2 hours post-induction of sporulation in cells expressing SpoIIE-3XFLAG or deleted for *spoIIE*. Yellow arrows indicate punctate DivIB along the cell membrane. *Pxyl-divIB* expression induced with 1 mM xylose.

**Figure S3:**
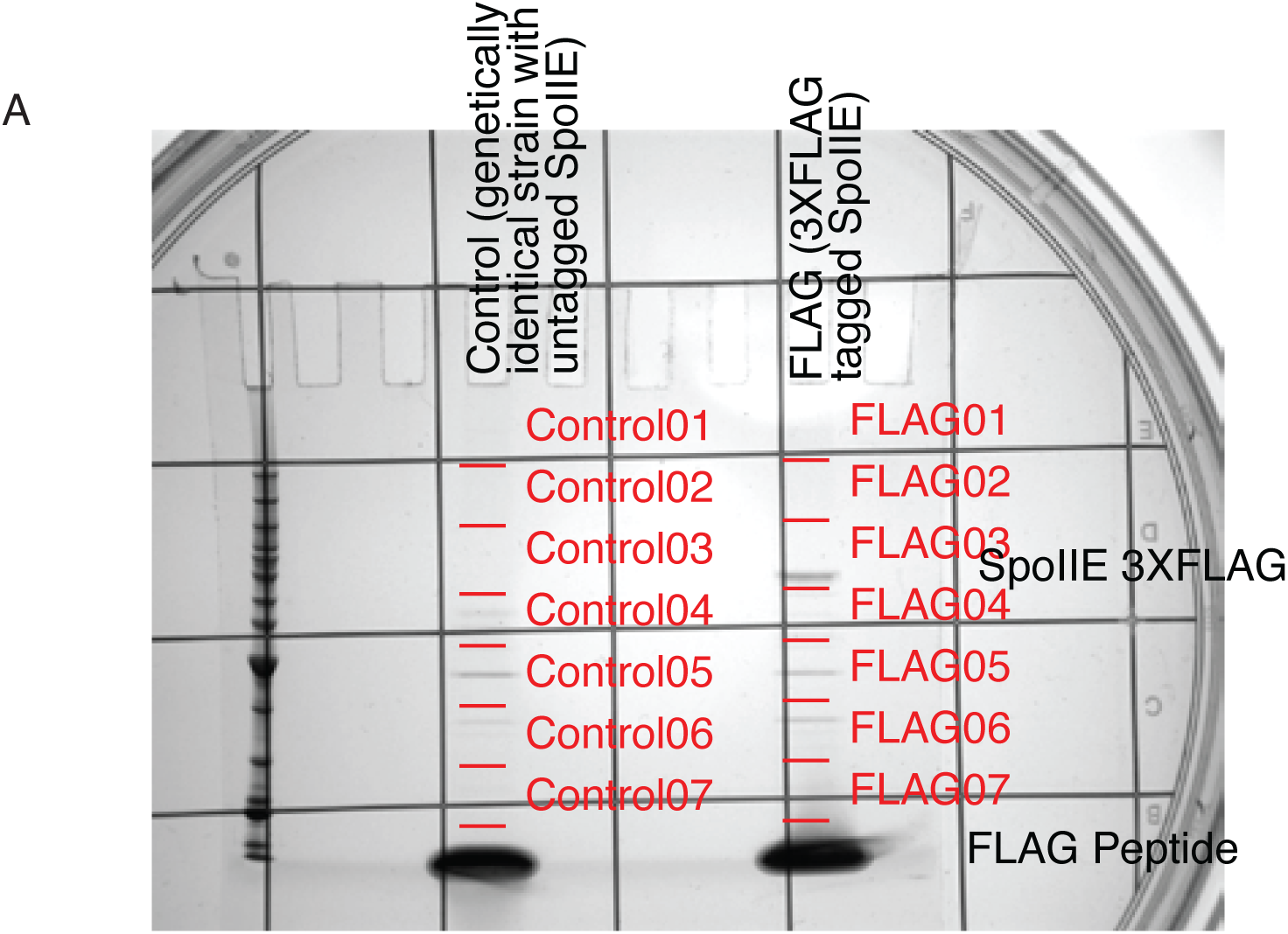
Identification of SpoIIE interaction partners by co-immunoprecipitation and mass-spectrometry. Shown is a gel of the co-immunoprecipitated samples from *spoIIE-3XFLAG* strain and a control strain expressing untagged SpoIIE. Each lane of the gel was cut into seven slices as indicated in the mage. Proteins were eluted from the gel, trypsinized, and subjected to mass spectrometry.

